# Protein function prediction in genomes: Critical assessment of coiled-coil predictions based on protein structure data

**DOI:** 10.1101/675025

**Authors:** Dominic Simm, Klas Hatje, Stephan Waack, Martin Kollmar

## Abstract

Coiled-coil regions were among the first protein motifs described structurally and theoretically. The beauty and simplicity of the motif gives hope to detecting coiled-coil regions with reasonable accuracy and precision in any protein sequence. Here, we re-evaluated the most commonly used coiled-coil prediction tools with respect to the most comprehensive reference data set available, the entire Protein Data Base (PDB), down to each amino acid and its secondary structure. Apart from the thirtyfold difference in number of predicted coiled-coils the tools strongly vary in their predictions, across structures and within structures. The evaluation of the false discovery rate and Matthews correlation coefficient, a widely used performance metric for imbalanced data sets, suggests that the tested tools have only limited applicability for large data sets. Coiled-coil predictions strongly impact the functional characterization of proteins, are used for functional genome annotation, and should therefore be supported and validated by additional information.

## Introduction

Coiled-coils consist of two or more α-helices that twist around each other and give rise to a multitude of supercoiled quaternary structures ^1,2^. Because of the simplicity of the smallest building block, two closely packing α-helices, whose coordinates can be calculated from parametric equations ^3^, the coiled-coil dimer was likely the first structural element, for which a sequence – structure – function relationship could be established ^4,5^. Accordingly, one of the first tools for predicting protein function was COILS, which allowed the identification of coiled-coil regions from protein sequences alone ^6^. Coiled-coil structures are claimed to be better understood than those of any other fold ^7,8^ and are increasingly used as building blocks in the emerging fields of synthetic biology and *de novo* protein design ^9,10^. The most advanced design case so far is likely a coiled-coil that can switch between pentameric and hexameric states upon pH-change ^11^. While writing up amino acid sequences forming coiled-coils with dedicated oligomeric state is possible now, the complementary approach to detect coiled-coil regions in amino acid sequences is likely to be resolved as well given the biochemical understanding of this structural motif. Even if the oligomeric state were not always predicted correctly, the presence of a coiled-coil should be noticed. Predicting the oligomeric state is indeed still error prone because of the low number of available protein structures with complex coiled-coil arrangements for training the algorithms ^12–14^. Accordingly, there are multiple studies indicating high sensitivity and specificity of the most current software packages ^12,15,16^. The benchmark data sets used, however, are limited by restriction to a selection of SCOP protein families containing coiled-coil regions (SCOP = Structural Classification of Proteins database; ^17,18^), or intersections of SCOP and SOCKET hits (SOCKET = tool to detect knobs-into-hole packing in protein structures; ^19^). This approach only allows assessing the sensitivity and specificity against highly restricted datasets and therefore strongly overestimates the proportion of true positive coiled-coil predictions. It does not allow to even estimate the number of false negative cases (no prediction where a coiled-coil is present) and false positive predictions (prediction of a coiled-coil where there is none) in a representative proteome.

Coiled-coil predictions are part of the standard tool box for the functional genome annotation. To our knowledge, NCOILS (identical to COILS v.2) is by far the most widely used tool in these functional annotations, likely because of its processing speed and its ease of use not requiring any additional software library or protein profile database. SpiriCoil, in contrast, scores query sequences against protein profiles of the SUPERFAMILY database, which is thought to represent all proteins of known structure ^20^, and passes SUPERFAMILY’s coiled-coil assignments to the new sequence ^12^. SpiriCoil was used to predict coiled-coils in the proteomes of more than 1,200 sequenced genomes suggesting that 0.33 to 6.53% of a species’ proteins contain at least one coiled-coil region ^12^. Proteome-wide predictions with NCOILS are of the same order ^21^. Because most of the proteins predicted to contain coiled-coils do not belong to the protein families with known extended coiled-coil regions such as muscle myosin heavy chain and intermediate filament proteins we wondered how many of these proteins really contain true coiled-coil domains. The most promising possibility to evaluate coiled-coil predictions is the comparison with their protein structures. Therefore, we assessed the current status of coiled-coil prediction accuracy by running all available coiled-coil prediction tools against all sequences, for which protein structures are known: the entire PDB. Each software was used with default parameters as recommended by the developers and as commonly done in genome annotation pipelines.

## Results

### Prediction of coiled-coils in protein structures and their sequences

To create a ground truth to rely on and compare against, we used SOCKET ^19^, the *de facto* standard to detect coiled-coil regions within PDB structures. The number of coiled-coils that SOCKET might miss is expected to be relatively small compared to the size of the PDB. Such cases could be NMR structures where SOCKET can have trouble dealing with ^12^. Within 144,270 PDB files (PDB status 12/2018), SOCKET detected 59,693 components (27,803 coiled-coils) in 10,684 (7.41%) PDB files (Figure 1A), which we defined as reference set (true positives). This means that most PDB files contain multiple coiled-coil domains, either in molecules related by non-crystallographic symmetry, in different regions of the same molecule, or in different molecules. The total number of sequences in the PDB files is 187,021 unique sequences (excluding all sequences related by non-crystallographic symmetry), which we left unreduced in terms of similarity or other criteria in order to have a broad, representative dataset. Every filter would introduce a bias on the data set or just subsample the data without any effect on the statistics. At the level of these 187,021 sequences, the 14,117 (7,51 %) distinct coiled-coils are similarly represented as in genome annotations ^12^. To assess the accuracy of coiled-coil predictions from sequences alone, we compared the SOCKET reference set with the results from NCOILS (COILS2.2; ^6^), PairCoil ^22^, PairCoil2 ^23^, MultiCoil ^24^, MultiCoil2 ^25^, and Marcoil ^26^. We did not include the tools SOSUIcoil ^27^ and RFcoil ^28^, because these are not accessible anymore. The tools CCHMM ^29^ and CCHMM-PROF ^30^ were also excluded, because these can only be accessed via a web interface, which is not applicable for a systematic evaluation. Furthermore, a short test for prediction performance using the globular and coiled-coil free myosin motor domain resulted in many predicted coiled-coil regions, which are obviously not correct (Supplementary Figure 1). SpiriCoil is in general only available via a web interface ^12^, caused internal server errors when used, and was therefore excluded. We also refrained from using the tool PCOILS ^31^, the successor of NCOILS, because it runs very slowly and is thus not applicable for the amount of data to be analysed. The latter problem is likely the reason why NCOILS is the tool commonly used in genome annotation projects. In addition, PCOILS was found to be less accurate than Marcoil ^15^. To best simulate a common functional protein annotation, we used default parameters for each tool as recommended by the developers, except for setting 21 amino acids as sliding window in all tools for comparability (21 is default in NCOILS; Figure 1).

**Figure 1:**
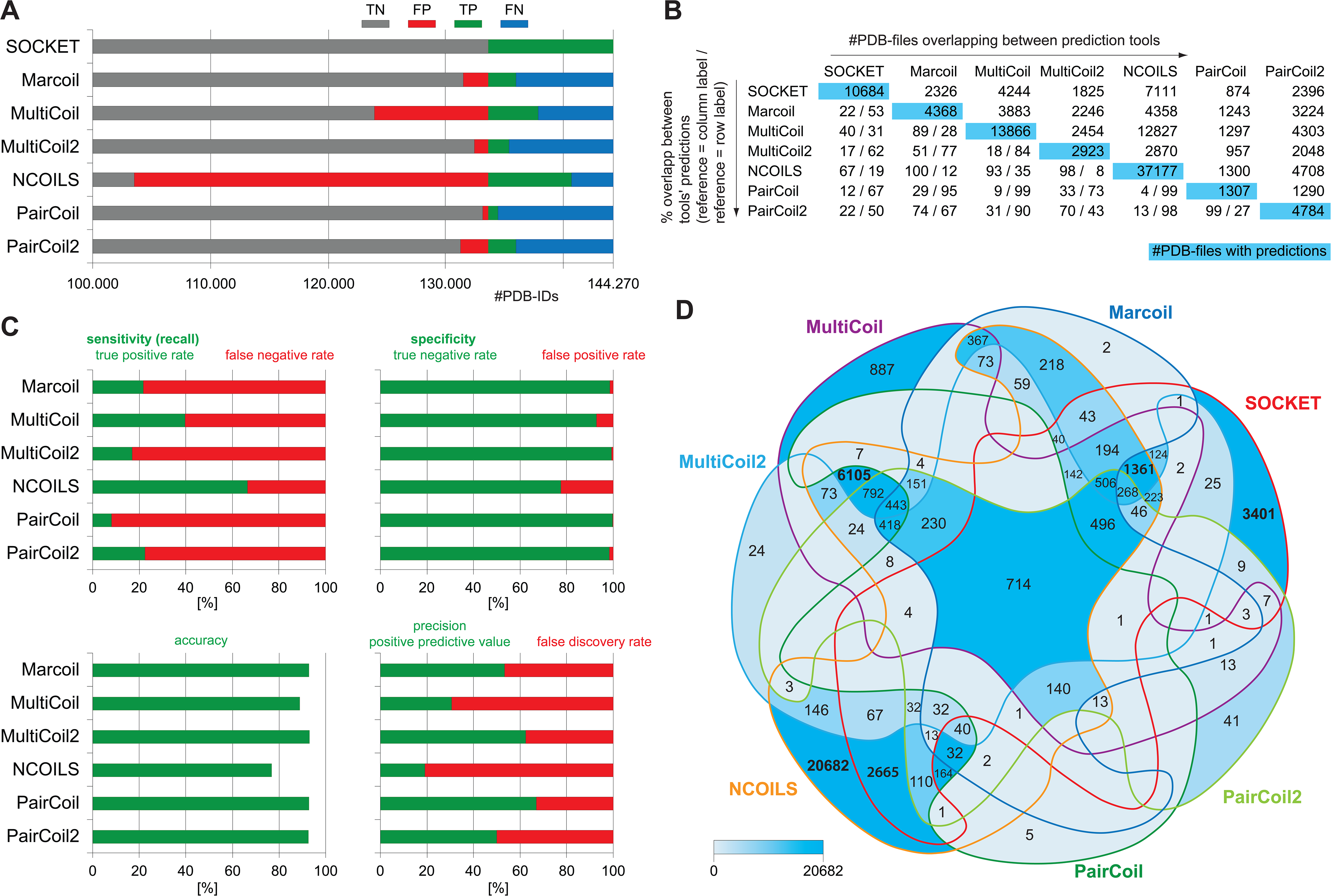
Coiled-coil regions identified by SOCKET and coiled-coil prediction tools at the level of PDB files. A) Of the 144,270 PDB files 10,684 contain a coiled-coil according to SOCKET (TP = true positives; the remaining 133,586 PDB files represent the true negatives=TN). B) The tools predicted coiled-coils in 1,307 (PairCoil) to 37,177 PDB files (NCOILS). C) Commonly used statistics for evaluating the performance of the coiled-coil prediction tools. Because of the characteristics of the benchmark data set, the specificity and accuracy of the tools are very high, while sensitivity and precision are rather low. It is obvious that the performance of the tools cannot be evaluated just based on these metrics. D) The 7-way Venn diagram shows the subsets of PDB files with predicted coiled-coils found by the respective combinations of tools colored by number in intersection. The intersection of PDB files with coiled-coils predicted by all tools is 714 PDB files, and 1,210 PDB files when ignoring PairCoil, the tool with the least predictions.

### Comparing coiled-coil predictions across PDB files

The various tools predict coiled-coils in strikingly different numbers of PDB files with PairCoil predicting coiled-coils in fewest (1,307) and NCOILS in most PDB files (37,177; Figures 1B). Each tool’s predictions overlap with 8.18% (PairCoil) to 39.72% (MultiCoil) of the PDB files with SOCKET hits (sensitivity; Figures 1B and 1C) showing that the tools did not predict any coiled-coil regions in the vast majority of the PDB files where SOCKET identified coiled-coils, except for NCOILS (overlaps with 66.56% of SOCKET hits). On the other side, 33.13% (PairCoils) to 80.87% (NCOILS) of the tools’s predictions were found in PDB files where SOCKET did not find any hit (false discovery rate; Figure 1C). Ignoring PairCoil, with which by far the fewest coiled-coils were predicted, the minimum overlap of PDB files with coiled-coil predictions from any two tools is 2,048 (Figure 1B). Although these numbers suggest considerable overlap of predictions in the same sequences, the opposite is found (Figure 1D). The intersection of PDB files with SOCKET hits and coiled-coils predicted by all tools is 714 PDB files. If ignoring PairCoil, the tool with the fewest predictions, this number increases to 1,210 PDB files. SOCKET hits are exclusive in 3,401 PDB files (31.8% of all SOCKET hits). All tools predicted coiled-coils in 230 PDB files (648 when ignoring PairCoil) where SOCKET did not identify any, potentially indicating structures outside SOCKETs default cut-off and structures difficult to resolve by SOCKET (e.g. some NMR structures). Coiled-coils predicted by at least two tools but no SOCKET hit were found in 9,399 PDB files, a considerably higher number than that from SOCKET hits overlapping with at least one of the tools (7,283 PDB files). While almost all PairCoil and Marcoil predictions overlapped with at least SOCKET or one other tool, PairCoil2, MultiCoil, MultiCoil2, and NCOILS exclusively predicted coiled-coil regions in 41 (0.86% of PairCoil2 predictions), 887 (6.40%), 24 (0.82%), and 20,682 (55.63%) PDB files, respectively (Figure 1D).

### Extent of structural overlap between coiled-coil predictions and SOCKET hits

Given already this diversity of PDB files with coiled-coil predictions and SOCKET hit regions the predictions could actually overlap with SOCKET hit regions or map to different regions on the same structures or different molecules within the PDB files. This aspect has been ignored in all comparative analyses so far and means that at the amino acid level the number of true positive predictions is even lower and the number of false positive predictions is even higher. Considering possible tool-dependent bias in determining start and end positions of the predictions we determined the number of predictions overlapping SOCKET hits in dependence of reference (either SOCKET or tool) and degree of overlap (Figure 2 and Supplementary Figure S2). Requiring at least a single amino acid overlap, only 26.66% (NCOILS) to 66.98% (MultiCoil2) of predicted coiled-coils overlap with SOCKET hits indicating that the majority of the predicted coiled-coils do not at all overlap with SOCKET hits, although SOCKET hits and predicted coiled-coils were found in the same PDB file. Taking the prediction tool as reference, only 13.43% (MultiCoil) to 34.02% (PairCoil) of the tools’ predictions overlap with at least 50% of their predicted sequence regions with the SOCKET hit regions. If requiring at least 80% overlap between predicted coiled-coil and SOCKET hit region, the proportion of overlapping regions decreases to 5.35% (NCOILS) to 17.48% (PairCoil). When taking SOCKET hits as reference, 23.56% (NCOILS) to 65.02% (MultiCoil2) of the predictions overlap with at least 50% of respective SOCKET hit regions. The percentages of overlapped SOCKET hit regions only slightly decrease with increasing size of overlap. This shows that predicted coiled-coil regions are in general considerably longer than SOCKET hit regions and therefore extend to protein structural regions not folded into knobs-into-hole packings.

**Figure 2:**
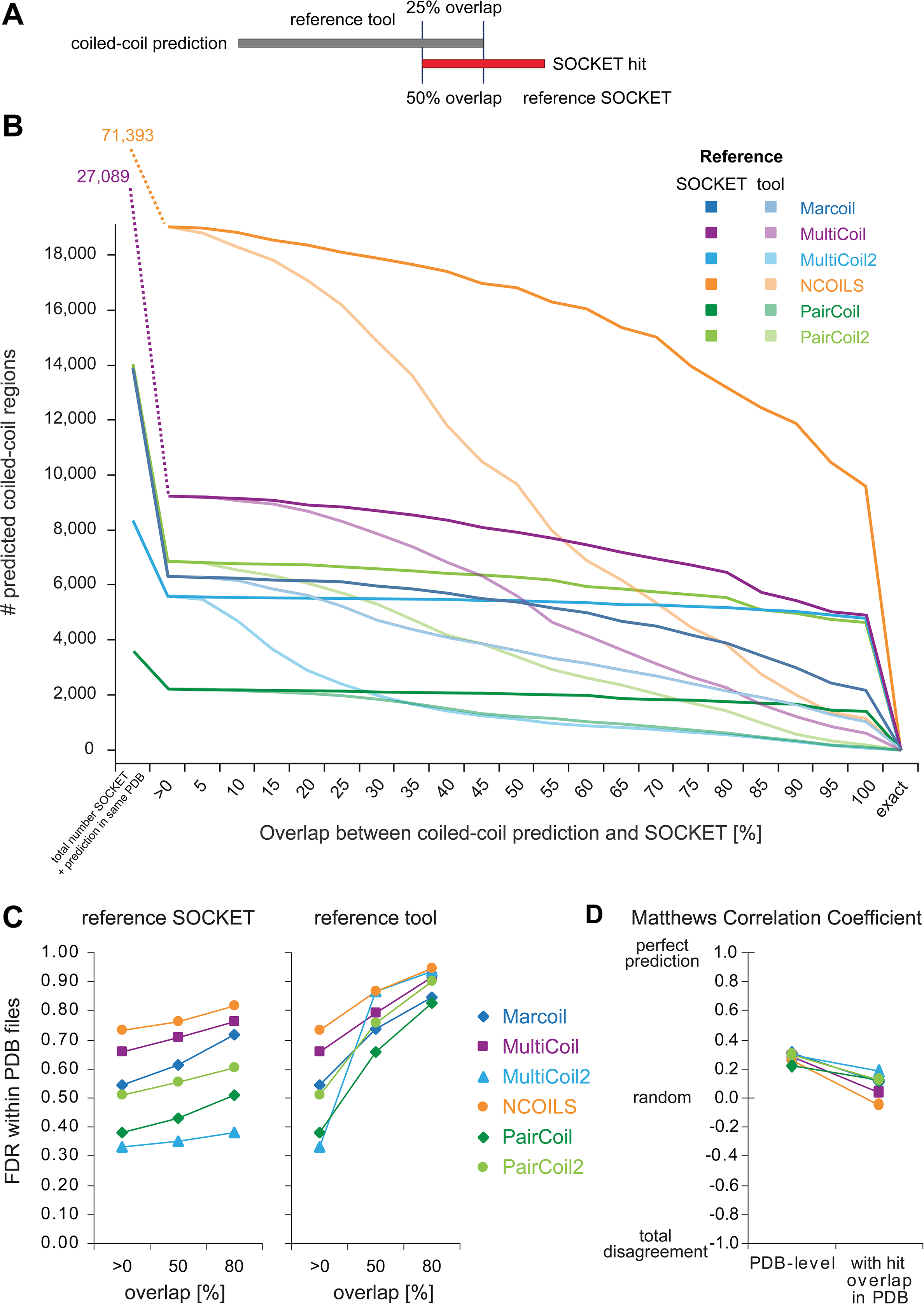
Overlap of coiled-coil predictions with SOCKET hit regions. A) Schematic drawing of a coiled-coil prediction overlapping a SOCKET hit. Depending on taking the tool or SOCKET as reference the proportion of the overlap is different. B) The starting point for this analysis was all PDB files with at least a SOCKET hit and a coiled-coil prediction independent of whether these were in the same sequence or overlapping (values at the left side). The percentage overlap varies in dependence of whether the SOCKET hit or the coiled-coil prediction is taken as reference. Plots based on the percentage of regions with overlap in the given range (in addition to the plot based on total numbers shown here) are shown in Supplementary Figure S2. C) False discovery rate when considering only those PDB files in which both SOCKET hits and coiled-coil predictions were found. Only those predictions were regarded as true positives that overlap with at least a single amino acid or at least 50% or 80% of their regions. To account for the different lengths of SOCKET hits and predicted coiled-coil regions, the overlap was evaluated separately by taking either SOCKET or prediction as reference. D) Matthews correlation coefficient at the PDB level (SOCKET hit and prediction in same PDB file) and the level of overlapping hits (SOCKET hit and prediction overlap by at least a single amino acid). A MCC of +1 indicates a perfect prediction, predictions with MCCs around 0 are no better than random, and a MCC of −1 represents total disagreement between prediction and reference.

When evaluating the real overlap of predictions with coiled-coils in the structures, the false discovery rates (FDR; the percentage of predicted coiled-coils where there are none) within PDB files range between 33.02% (minimum a single amino acid overlap) and, when requiring 80% overlap, 81.51% when taking the SOCKET hits as reference for overlap, and 94.62% when taking the predicted coiled-coils as reference for overlap (Figure 2C). These false predictions within PDB files have to be seen in addition to the 33.13% to 80.87% false predictions determined already at the PDB file level. All described metrics measure the classification quality of either of true and false positives and negatives, and all rely on the specifics of the data set, i.e. the proportion of true positives and negatives. In contrast, the Matthews correlation coefficient (MCC) is a balanced measure of all classes returning a value between −1 and +1 ^32^. At the PDB file level (SOCKET hit and coiled-coil prediction in same PDB file) the MCCs of the evaluated prediction tools range between 0.22 (PairCoil) and 0.31 (Marcoil) indicating that these predictions are only slightly better than random (Figure 2D). When requiring overlap of at least a single amino acid between SOCKET hit and coiled-coil prediction, the MCCs decrease to −0.05 (NCOILS) to 0.19 (MultiCoil2).

### Patterns of coiled-coil predictions

Coiled-coil regions are characterised by hydrophobic residues at the interface between the supercoiled α-helices and by charged and polar amino acids at the outside. This pattern is usually found in heptads with the corresponding amino acids marked as *abcdefg* although slightly different patterns from hendecad and pentadecad repeats are observed ^33,34^. The expected pattern is found in all SOCKET regions, those which overlap regions where tools also predict coiled-coils, and those where tools did not identify any (Figure 3 and Supplementary Figure S3). There is a clear preference for leucines in *d* positions and increased propensity for isoleucines and valines in *a* positions compared to *d* positions. There is strong discrimination against glutamates in *a* positions and lysines and arginines in both *a* and *d* positions. However, there are significantly more leucines in *e* and *g* positions than in *b, c*, and *f* positions, and there is no discrimination against glutamates in *d* positions (Figure 3). The most obvious difference between the patterns of only SOCKET hits and those with overlapping predictions is the slightly lower preference for and discrimination against leucines and charged residues, respectively. The comparison indicates that all the SOCKET-determined coiled-coils in the 3,401 PDB files, where the tools did not predict any coiled-coil, are not very different from the other SOCKET hits from the perspective of amino acid distribution at heptad positions. The patterns of the predicted coiled-coil regions are very similar with respect to each other and to the SOCKET pattern although more discriminating against glutamates and lysines in *a* and *d* positions (Figure 3).

**Figure 3:**
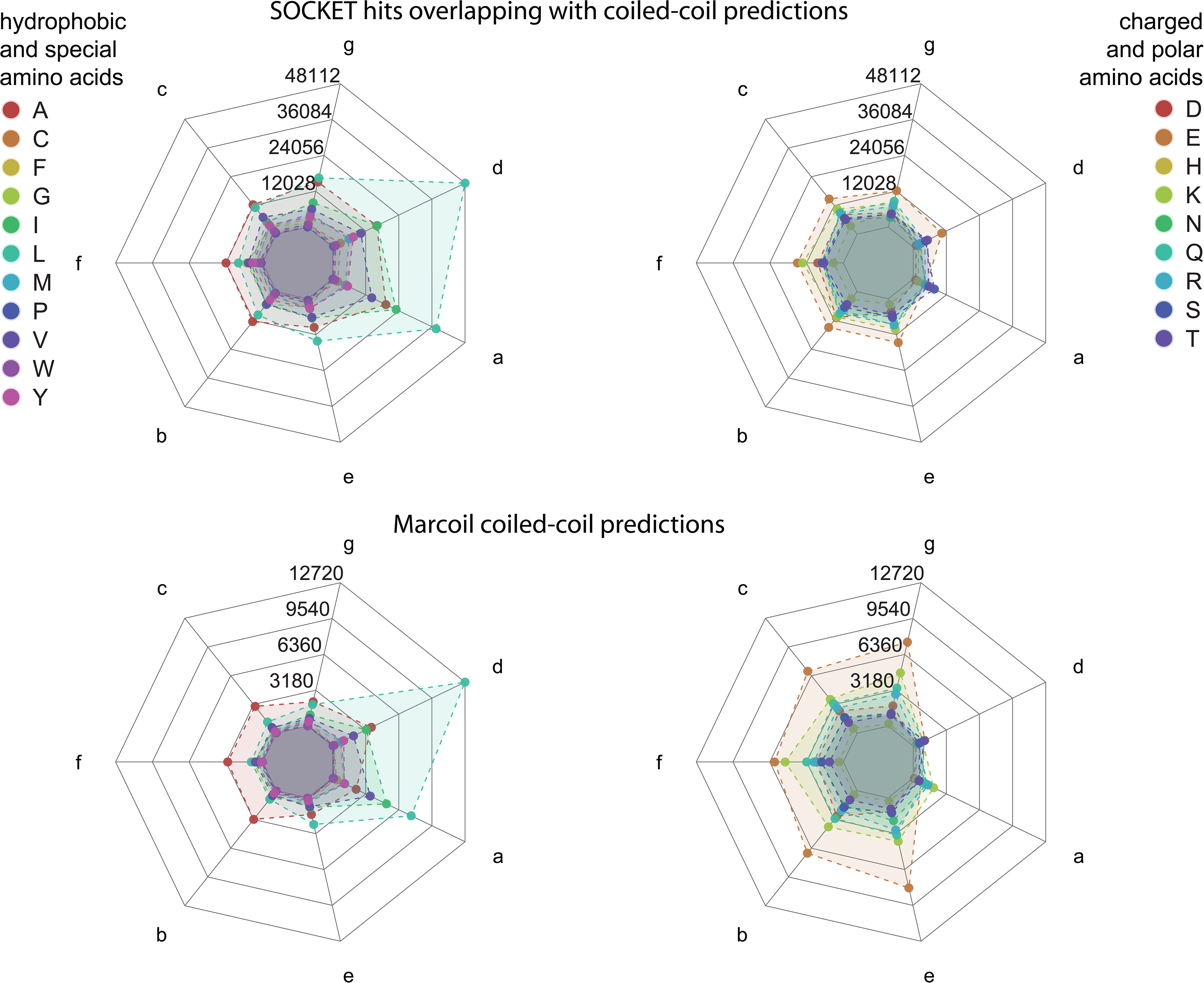
Amino acid preferences at heptad positions *abcdefg*. It is well known that hydrophobic amino acids are preferred at the interface between coiled α-helices, at positions *a* and *d*, and that charged and polar amino acids are preferred at the outside, especially at positions *e* and *g*. Oppositely charged amino acids at positions *e* and *g* are thought to further contribute to coiled-coil stability through charged interactions. Because SOCKET not only detects “classical” coiled-coils but interacting α-helices within globular protein structures, the distribution of hydrophobic and charged amino acids is slightly less biased in the latter structures (Supplementary Figure 3). Marcoil predictions show strong bias for leucine and isoleucine at the interior positions *a* and *d*, and for glutamate at all other positions. The heptad patterns of the other predicted coiled-coils show similar distributions. The letters at the axes denote the heptad register positions. Data values at grid lines refer to amino acid counts at each heptad position over all heptads.

The different curves for SOCKET hits overlapping predictions and predictions overlapping SOCKET hits (Figure 2) already showed that coiled-coil predictions are in general longer regions than SOCKET hits. This is supported by the length distributions of the coiled-coil regions (Figure 4). Most SOCKET hits are 10 to 19 amino acids long, while most Marcoil, MultiCoil and NCOILS regions are 20 to 29 amino acids long, and most PairCoil and PairCoil2 regions are 30 to 39 residues long. MultiCoil2 regions show a very different length distribution with no specific preference for a certain length. Instead, MultiCoil2 seems to combine consecutive helices, e.g. the repeat regions of helical bundle forming proteins such as spectrin and α-actinin, into single super-long coiled-coil regions.

**Figure 4:**
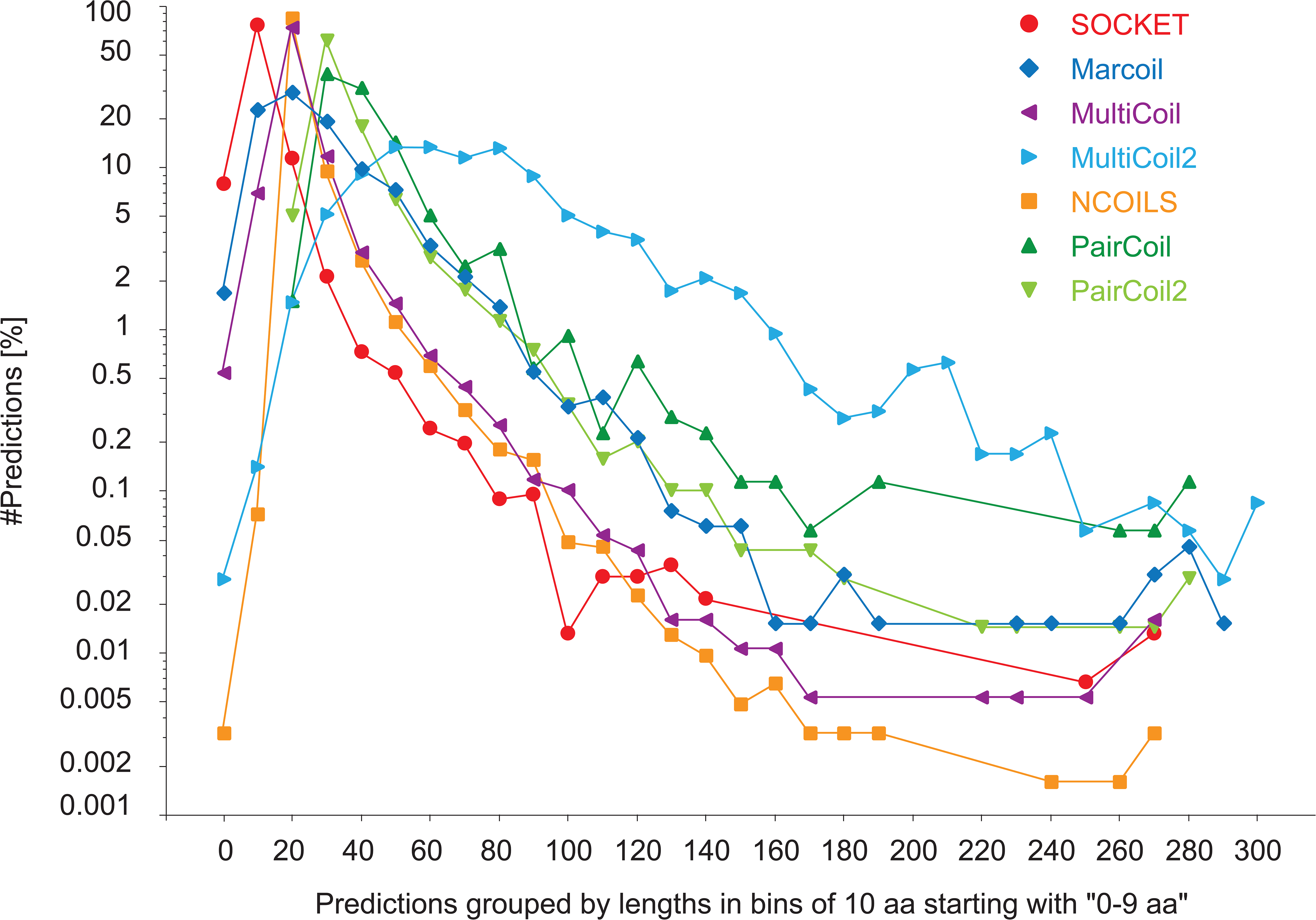
Length of SOCKET hits and coiled-coil predictions. SOCKET hits and coiled-coil predictions were grouped by length in bins of ten amino acids, and the number of hits/predictions in each bin plotted in percent with respect to the total number of hits/predictions of the respective tool.

### Coiled-coil predictions map to all types of secondary structural elements

The considerably deviating matchings of SOCKET-determined coiled-coil regions and predicted coiled-coils with respect to the PDB structures prompted us to look at the secondary structural elements of matched regions. As ground truth for the secondary structure we relied on the DSSP assignment ^35,36^. As expected by SOCKET’s algorithm, 99.994% of the amino acids within SOCKET hits match to α-helical regions (H in DSSP notation) while the remaining 0.006% match to loops (“blank“; Figure 5A). In contrast 20.78% (Marcoil) to 31.54% (NCOILS) of the regions predicted to be coiled-coils do not fold into α-helices, and considerable parts of MultiCoil (2.64%), MultiCoil2 (2.60%), and NCOILS (4.96%) predictions match to β-strands (Figure 5A). To allow visual inspection of these rather surprising results and detection of potential mis-assignments or systematic deviations we implemented a web-interface to the analysis database providing a search interface, a structure viewer, and sequence-based representations of all predictions in comparison. The web-interface can freely be accessed at https://waggawagga.motorprotein.de/pdbccviewer.

**Figure 5:**
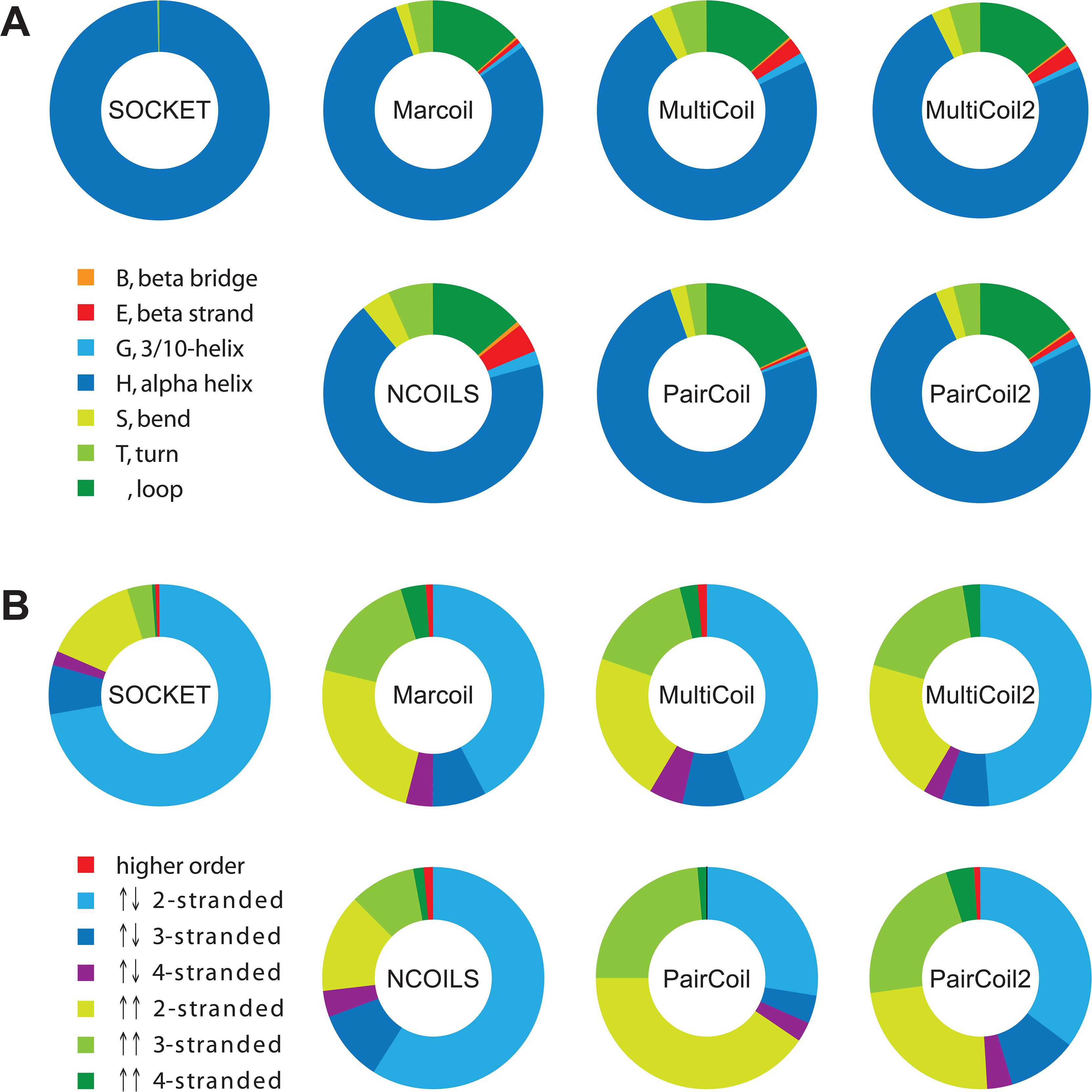
SOCKET hits and coiled-coil predictions matching protein structures. A) The plots represent the matching of all SOCKET hits and all coiled-coil predictions with secondary structure elements as determined by DSSP. The secondary structure assignment for each amino acid was read from the DSSP output, the assignments summed up for each element, and the distribution of elements determined for each tool in percent. B) Coiled-coil predictions on sequence alone are not biased for certain oligomeric states. As reference, the distribution of oligomeric states as determined by SOCKET is shown, with parallel and antiparallel arrangement of the α-helices separated. For each coiled-coil prediction tool, only those predictions were selected that overlap at least 50% of a SOCKET hit, and the oligomeric state assignments of the respective matched hits were collected.

### Parallel and antiparallel coiled-coils, and oligomeric states

Coiled-coils can have parallel and antiparallel arrangements of the α-helices and take part in many different oligomeric assemblies ^37^. The “classical” coiled-coil is a parallel homodimer, and accordingly the protein training data of the first prediction tools consist of parallel homo- and heterodimers such as myosins, tropomyosins, kinesins, and intermediate filament proteins. In the PDB by far the most detected arrangement is the antiparallel 2-stranded coiled-coil (72.1%) followed by the parallel 2-stranded coiled-coil (14.1%; Figure 5B).NCOILS’ predictions have the largest overlap with SOCKET hits and thus the most similar distribution of arrangements with SOCKET. PairCoil shows the strongest bias of all prediction tools for detecting parallel 2-stranded coiled-coils. Marcoil, MultiCoil, MultiCoil2, and PairCoil2 all have similar distributions with bias towards parallel 2-stranded and 3-stranded coiled-coils (Figure 5B). These data show that the prediction algorithms do not exclude certain arrangements, be it the direction or the number of involved α-helices.

## Discussion

Coiled-coils consist of a minimum building block of just an α-helix and can be designed on a drawing board based on a poly-alanine backbone and subsequently substituting alanines by hydrophobic, charged, and polar amino acids to obtain structures with certain characteristics, mainly a certain length and topology ^9,10^. The simplicity in design should, in principle, allow a relatively accurate and precise prediction of these motifs in real-world sequences. For the evaluation of the performance of coiled-coil predictions in the context of a functional genome annotation the reference data set should be large and diverse, should contain only a few percent sequences with coiled-coil regions, and should allow structural verification. To our knowledge the protein structure databank (PDB) represents the most comprehensive reference data set given the broad sampling of species, protein families and protein folds. As ground truth and reference we used the coiled-coils detected by SOCKET, which identifies knobs-into-hole packings of α-helices within protein structures. We evaluated the performance of coiled-coil prediction tools against all sequences within the PDB using all common binary classification metrics. The specificity and accuracy of all prediction tools is very high, which is a natural result from the large proportion of true negatives within the data set. In contrast, the sensitivity and positive predictive value are rather low. This is, in part, due to the lower number of predictions than SOCKET hits in general (however, NCOILS predicted about four times more coiled-coils), but more importantly the result of the large proportion of false positive predictions. Coiled-coils were predicted in 31,040 PDB files where no SOCKET hits were found, and in addition thousands of coiled-coils were detected within PDB files that do not overlap with SOCKET hits. The problems that SOCKET has with some structures might explain some dozens of these predicted coiled-coils and even a few hundreds, but not the majority. It is highly unlikely that SOCKET missed many “classical coiled-coils”, which are the supposed primary target of the prediction tools. Without having inspected all predictions manually, we suspect that it is more likely that most of these predictions are in fact false positive hits. These include the reported coiled-coil predictions in polyQ regions (see Supplementary Notes for more details).

Because of the low sensitivity the false negative rates are high. In 3,401 PDB files coiled-coils were exclusively found by SOCKET. At first instance, the most likely explanation for these cases is that the coiled-coil prediction tools are thought to be specific for solvent-exposed, left-handed coiled-coil dimers, and are not expected to detect types of coiled-coil α-helices buried within globular domains or as part of transmembrane structures. And because most coiled-coil prediction tools were developed before next-generation sequencing boosted sequence databases, the relatively low number of training data for tool development could have also been limiting in detecting more divergent coiled-coil types. However, the false negative rates of the individual tools at the PDB file level are in the range of 60.28% (MultiCoil) to 91.82% (PairCoil) with the exception of NCOILS (33.44% false negative rate). In addition, the amino acid patterns at the heptad positions of SOCKET hits overlapping and not overlapping with predictions are very similar. This indicates that most of the SOCKET hits in the 3,401 PDB files could just as well have been identified by coiled-coil prediction tools as those that were detected. While the discussed metrics depend on the proportion of true and false positives and negatives in the benchmark data set, the Matthews correlation coefficient is insensitive to the proportion of coiled-coils in the data set and gives a more balanced assessment of the performance. The performance at the PDB file level was rather poor (MCCs between 0.22 and 0.31) and significantly decreased further when requiring only the overlap of a single amino acid between SOCKET hits and coiled-coil predictions within PDB files (MCCs between −0.05 and 0.19). The MCCs do not considerably increase if the 3,401 PDB files with exclusive SOCKET hits are regarded as true negatives (no coiled-coils to be detected by prediction tools) and if the 230 PDB files with coiled-coils predicted by all tools but not detected by SOCKET are regarded as true positives. Requiring only a single amino acid overlap is a rather weak criterion and balances possible biasing effects from software parameters such as the SOCKET packing-cutoff and window sizes or cutoffs from prediction tools.

The finding that coiled-coil prediction tools show low performance when benchmarked with sequences from heterogeneous protein structures is not completely new. A comparison of SpiriCoil, Marcoil, and PairCoil2, revealed a similar low absolute performance, with SpiriCoil, the supposed best performing coiled-coil prediction tool in this comparison, displaying a sensitivity of 41.7% and an FDR of 84.6% at the level of sequences ^12^. In this comparison, 2.71% of the sequences in the test data contained coiled-coils, which is slightly lower than the percentage of likely coiled-coil regions in our data set (7.41% of 144,270 PDB files contained SOCKET hits). However, SpiriCoil just passes the coiled-coil assignment from SUPERFAMILY protein profiles on query sequences based on global sequence comparisons without ever verifying the presence of a coiled-coil. This approach therefore ignores domain gain, loss, and rearrangement processes, which are very common in eukaryotic genomes. In contrast, the latest comparison showing good sensitivity and specificity of the prediction tools was based on a highly biased sequence data set with 63.4% of the 1643 test sequences containing coiled-coils, and each of these sequences containing 2.09 coiled-coil regions on average ^16^. Already an even and random assignment of coiled-coils to sequences of this data set would result in sensitivity and specificity of 79% and 100%, respectively. In addition to these general shortcomings in approach and data set, in both previous studies the precise location of the coiled-coils with respect to reference SOCKET hits and secondary structural elements was not determined.

Given this analysis and the application of the evaluated prediction tools, especially NCOILS, in the functional annotation of genomes it is highly questionable that many of the proteins with predicted regions really contain coiled-coil domains. In addition it is highly likely that coiled-coil domains have been missed in many proteins. Given the broad application of coiled-coil prediction tools, as citation rates suggest, and the high interest in this structural motif, as publication numbers suggest, we see a high demand for accurate coiled-coil prediction. We suggest improving the tool’s performance against unbiased and not pre-selected data and to use approaches that combine sequence profiles and secondary structure assignments, or that discriminate against certain atypical features. Secondary structure information, for example, was included in WDSP, a pipeline to predict WD40 repeats and domains ^38^. WD40 repeats have very low sequence homology, are therefore notoriously difficult to detect, and are usually present in a chain of seven to fold into a domain ^39,40^. In WDSP, protein sequences are filtered selecting fragments with β-sheets according to PSIPRED ^41^, subsequently WD40 repeats are detected using a profile generated by aligning repeats by secondary structure elements and not global similarity, and finally WD40 domains are assigned when chains of at least six WD40 repeats are present. As another example, Waggawagga uses the discriminative approach to detect stable single α-helices (SAH domains), which coiled-coil prediction tools mis-predict as coiled-coils ^42–44^. Waggawagga searches for networks of oppositely charged residues and discriminates against helix-breaking residues, networks of residues with identical charge, and networks of hydrophobic residues as found in the hydrophobic seams of coiled-coils. Approaches similar to those used by WDSP and Waggawagga could be implemented to improve coiled-coil predictions. For example, protein sequence regions could be pre-filtered and/or coiled-coil predictions could be post-filtered by secondary structure predictions. A selection filter for potentially coiled-coil domain containing regions could also be the detection by at least two tools. Training the prediction tools against unbiased data such as the entire PDB could also improve tools’ performance. The evaluation of multiple tools to predict the pathogenicity of SNPs ^45^ and protein stability ^46^ also demonstrated low performance (low to medium MCCs), but the results stimulated substantial tool improvement with respect to the benchmark data sets.

In conclusion, at best, the evaluated tools predict coiled-coil regions in well described and well analysed coiled-coil forming proteins with reasonable accuracy. For predicting coiled-coils in large data sets with balanced proportion of all protein folds, such as present in gene prediction datasets, the tested tools have only limited applicability. One possibility to reduce the number of false predictions in such functional genome annotations would be to only accept coiled-coils regions if predicted by multiple prediction tools and to only predict coiled-coils in regions not already covered by other protein domain predictions.

## Methods

### Benchmark dataset

To benchmark the performance of coiled-coil prediction software as fairly and reliable as possible, we created a copy set of the current state (15/12/2018) of all available 147,073 PDB structures from the RCSB Protein Data Bank ^47^. The flatfiles were downloaded, stored locally and parsed with BioRuby v.1.5.1 ^48^. Handling issues with some PDB-files reduced the number of usable structures to 144,270. Main reasons were unavailability (moved/renamed/discontinued?) at the RCSB servers (761 files) and BioRuby parsing issues with some of the structures in the new PDBx format, introduced in 2014, some early structures and structures with non-natural amino-acids. The information from PDB files and all additional data generated were stored in a PostgreSQL database. To facilitate data handling and analysis of the PDB, DSSP, SOCKET, and coiled-coil prediction information a relational database scheme was designed, which stores the relevant data for the evaluation with low redundancy and depicts each data type into its assigned classes (Supplementary Figure S4). Protein sequences were extracted from the ATOM records of the PDB files. Unfiltered, all sequences break down to 187,776 unique sequences. In order to create a broad, representative data set, the sequences stay unreduced in terms of similarity or other criteria, even very short sequences were left in the data set. Only the 755 sequences containing amino acids labeled “unknown”, one-letter code “X”, which are handled very differently by the coiled-coil prediction tools, were removed from the analysis. The remaining 187,021 sequences were used as reference for all analyses. Accordingly, the secondary structure for every single amino acid within the reference sequences is known. The overall size of the database for the current PDB structure data set amounts to 2.9 GB, the flatfiles in combination with the predictions sum up to around 159 GB in 2.8 million files.

### Running coiled-coil software

The SOCKET algorithm relies on available secondary structure assignments from DSSP ^35,36^, which we generated according to the software documentation. For the determination of coiled-coil regions SOCKET ^19^ was used with the recommended parameter settings, especially the packing-cutoff was left at the default 7 Å as described in the documentation. The coiled-coils were, according to the database model, split into their superordinate structure and building/participating components, which contain registers, sequences and position information. To prevent mis-assignment of any amino acid due to sequence gaps or other unexpected shifts, the register assigned sequence of each SOCKET-determined coiled-coil component is searched in the respective PDB sequence and a potential offset is added to the component database entry.

Each coiled-coil prediction software was run with its recommended default settings and a search window of 21 amino acids, except for Marcoil and MultiCoil2, which are implemented to run without a respective window setting. Accordingly, quite conservative thresholds were set for coiled-coil selections. For Marcoil a lower limit of 90.0 (minimum) was chosen (HMM training file 9FAM, with default transition and emission parameters). For MultiCoil and MultiCoil2, the “CoiledCoil-Threshold” was set to 0.25 (minimum). For NCOILS, the “CoiledCoil-Threshold” was 0.5 (minimum), and the latest provided MTIDK-matrix was used. For PairCoil and PairCoil2, the “CoiledCoil-Threshold” was 0.84 (minimum) and 0.025 (maximum), respectively. Because the reference sequences do not contain the gap information as present in protein structures, coiled-coil prediction tools handle all sequences as continuous entities. This might affect the length of some predicted coiled-coils.

### Determining overlap between assignments and predictions

Assigning SOCKET hits, DSSP assignments, and coiled-coil predictions globally to PDB files and chains is trivial. However, precisely determining overlapping regions within sequences is more challenging because amino acid numbering schemes change with data parsing and gaps in structures. Numbering of amino acids in DSSP features and SOCKET hits follows the numbering of the amino acids in the structures, which either start with the first amino acid of the protein construct, the first amino acid of the sequence of interest (excluding any terminal amino acids from protein expression plasmids), or follow the numbering of the analysed protein with respect to the numbering in the gene or transcript. The numbering of amino acids in the structures is usually aware of gaps. When extracting sequences from PDB files all position-wise numbering information including gaps is discarded, only start- and end-positions in PDB numbering are retained. Sequences themselves are stored plain without any numbering in the database, meaning a sequential numbering starting with “1” when referred to from other tools. Instead of fitting the results from the prediction tools to the complex numbering in the structure files, the numbering of the initial register sequences of the SOCKET hits was shifted to the position-independent numbering as described. Accordingly, the matching between SOCKET hits and coiled-coil predictions is independent of any peculiarities in structure numbering and independent of any gaps in the structures. Similarly, one after another the sequences corresponding to every contiguous DSSP feature in a structure are located in the number-less sequences and the DSSP features are subsequently numbered according to the matching. A problem with this approach could be that very short DSSP features (1-3 amino acids), which are surrounded by sequence without feature (“loop”), might match to more than one position. However, these very short DSSP features are very likely not part of SOCKET hits or coiled-coil predictions so that the matching of DSSP and SOCKET/coiled-coil prediction is not affected.

### Handling difficult cases

There are a few cases which cannot be resolved consistently without introducing multiple subcategories, which in turn would considerably detract from the main message without adding additional understanding. One of these problems is handling cases of overlapping SOCKET hits and coiled-coil prediction when one overlaps multiple of the other. In such cases we treated every overlap independently. For example, a long coiled-coil prediction could overlap with a SOCKET hit in its N-terminal half and another SOCKET hit in its C-terminal half. Such cases were treated as two independent overlap instances.

The actual data categories and assigned categories (true and false positives and negatives) for computing the Matthews correlation coefficient at the level of PDB files are clearly defined. It is, however, difficult to define the same categories (true and false positives and negatives) in case of evaluating the cases of overlapping SOCKET hits and coiled-coil predictions. The problem is that multiple hits were found in many PDB files, and those hits can be overlapping and non-overlapping. As a rough approximation for an upper bound we computed for each tool the percentage of overlapping hits within PDB files, and applied this percentage onto the number of overlapping PDB files. With this approach we rather overestimate the number of true overlapping hits, because for most tools there are more non-overlapping than overlapping hits in PDB files with both SOCKET hits and coiled-coil predictions, and the false positive PDB files contain, to some extent, multiple coiled-coil predictions whose contribution is ignored.

### Viewer for coiled-coils mapped to PDB structures

For visual inspection of SOCKET hits and coiled-coil predictions, a simplified search interface to the analysis database and a 3D molecule viewer were integrated into the coiled-coil project site Waggawagga. Structures can be searched by PDB ID and results are displayed for each structure on a single page. The result page presents the structure in JSmol, a JavaScript-only version of Jmol ^49^, for interactive viewing, some general information about the PDB file, and all SOCKET hits and coiled-coil predictions if found. The SOCKET hits and coiled-coil predictions are shown in the sequence-based and interactive Waggawagga format ^43^ providing access to all information down to the amino acid level. By simple selection SOCKET hits and coiled-coil predictions can be loaded and combined into the JSmol viewer. Intersecting regions between SOCKET hits and predictions are marked separately allowing the structure-based visual inspection of coiled-coil assignment by SOCKET versus prediction tools.

## Author Contributions

D.S. and M.K. designed the study with input from K.H.. D.S. developed the software, set up the data viewer, and performed data engineering and all computations. K.H. contributed to database design. D.S. and M.K. performed data analysis, designed the figures and drafted the manuscript. S.W. was involved in statistical analyses. All authors discussed the results and commented on the manuscript.

