## Supplementary Material for "Protein function prediction in genomes: Critical assessment of coiled-coil predictions based on protein structure data"

### Supplementary Notes

#### The polyglutamine puzzle

It has been suggested that (Q/N)-rich prions and polyQ-expanded proteins form coiled-coil structures based on the polyQ and neighbouring regions [1]. There are 18 structures now in the PDB containing stretches of at least six glutamines (Table S1). In only one of these, 1PJS, the polyQ region is part of a coiled-coil structure, while in the others these regions do not take part in any oligomerisation. In 1PJS the polyQ region is the extension of a very long α-helix. In other structures, the first two to four glutamines succeeding an extended α-helix often extend this α-helix while the remaining glutamines are mostly flexible and not visible in the crystal structures. Coiled-coil regions in polyQ proteins were detected with COILS and PairCoil2 [1]. However, NCOILS and also Marcoil already predict "coiled-coils" in artificial polyQ stretches with just any single other amino acid somewhere in the middle of the stretch. Similarly, these tools predict "coiled-coils" in artificial polyN stretches with a hydrophobic and a charged amino acid somewhere within the stretch. While polyQ and polyN regions might transiently fold into partially α-helical regions, it is highly unlikely that these cause specific protein interactions or form coiled-coils. Even if polyQ regions formed α-helices, the identical helix surface would lead to uncoordinated aggregation in all directions. However, organisms with massive homopolymeric regions such as *Dictyostelium discoideum* [2] do not show more protein aggregation than any other species. We suggest using the term "coiled-coil" in the original sense coined by Francis Crick for two or more α-helices in a dedicated structural packing, and not for any stretch of amino acids that might partially fold into α-helices and aggregate.

**Table S1**: Crystal structures of proteins containing stretches of at least six glutamines

| PDB id | polyQ stretch | observed structure |
|---|---|---|
| 1U6F | $QLQQLQ_6$ | turn, bend |
| 2DMS | $Q_7$ | turn, bend |
| 2OTU, 2OTW | $Q_{10}G$ | turn, bend, loop |
| 1QB3 | $Q_{16}HQTQ$ | no structure |
| 3IO4, 3IO6, 3IOR, 3IOT, 3IOU, 3IOV, 3IOW | $Q_{17}$ | multiple conformations, mainly loop and bend, partially helical |
| 3PJS | $QEQ_6$ | helical |
| 4A9Z | $Q_6$ | helical, turn |
| 4FE8, 4FEB, 4FEC, 4FED | $Q_7HQHQHQ_{27}$ | helical, loop, β-hairpin |

### Supplementary Figures

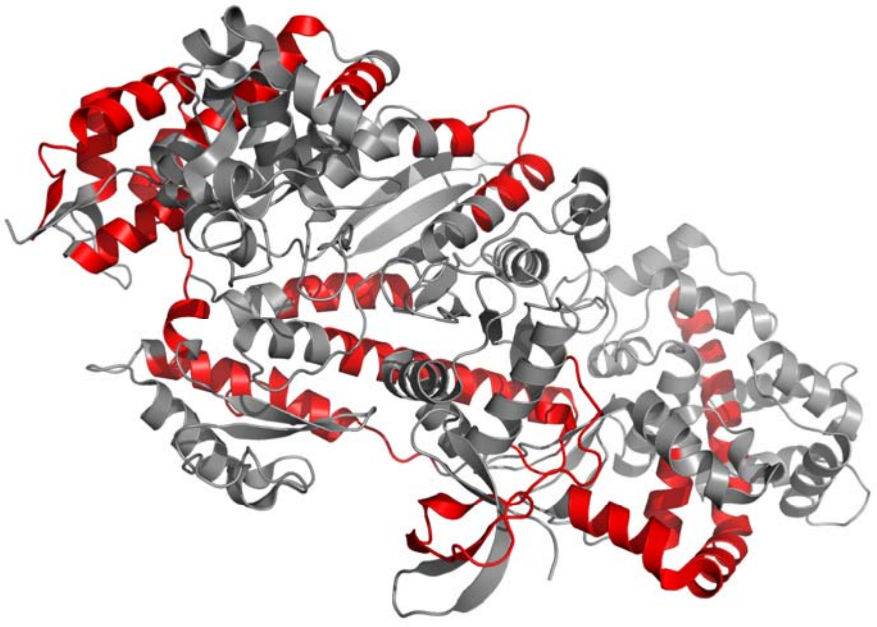

**Figure S1: Prediction of coiled-coils in human MyoVI with CCHMM-PROF.** To test the performance of CCHMM-PROF, which is only accessible via a web interface, the motor domain sequence of human MyoVI was used. MyoVI is a backwards walking myosin motor, and was chosen by chance from the list of available protein crystal structures of myosin motor domains. Predicted coiled-coil regions were mapped onto the crystal structure, PDB ID 2BKH,

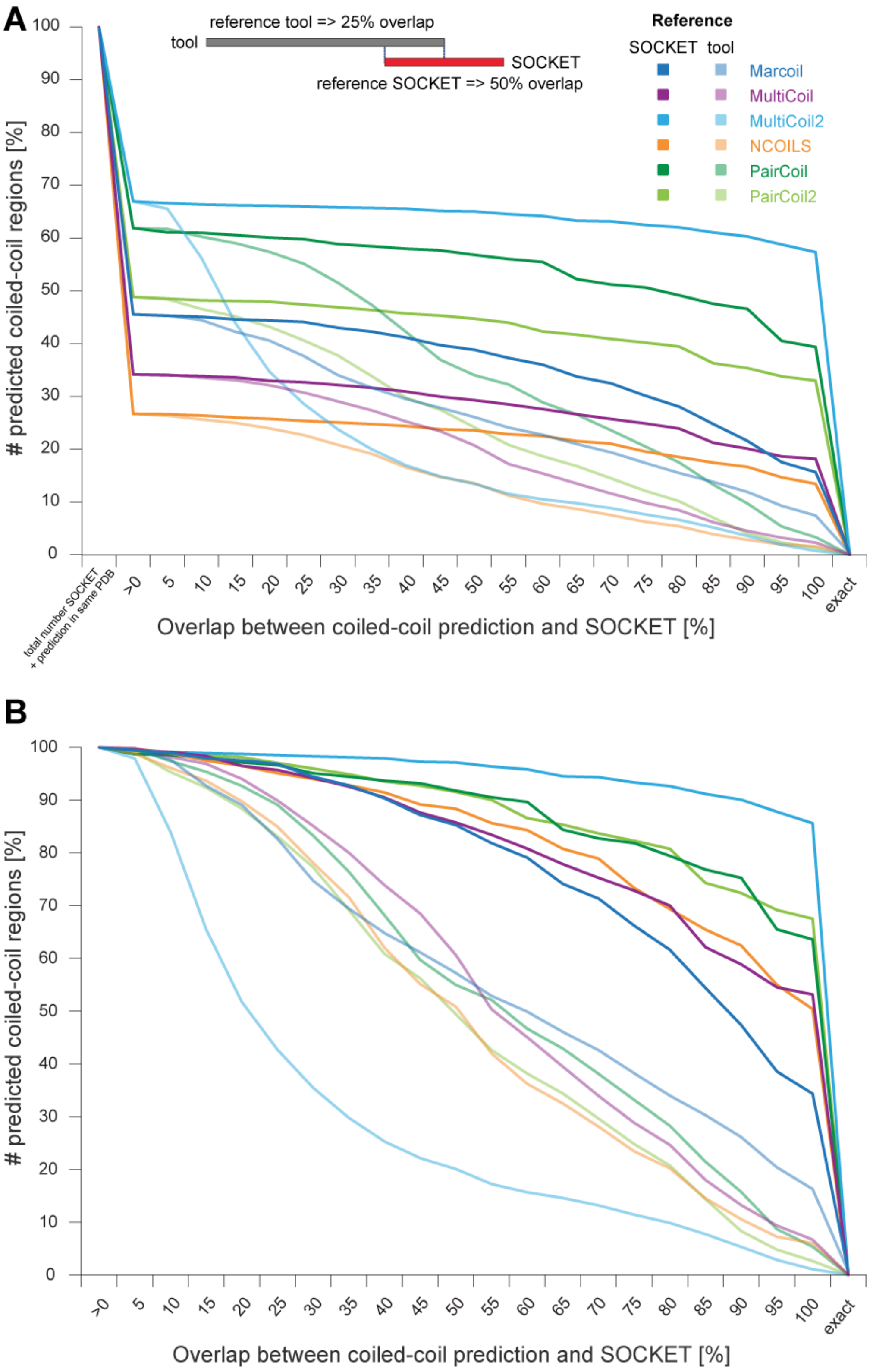


**Figure S2: Overlap of coiled-coil predictions with SOCKET hit regions.** The starting point for this analysis was all PDB files with at least a SOCKET hit and a coiled-coil prediction independent of whether these were in the same sequence or overlapping (values at the left side). The percentage overlap varies in dependence of whether the SOCKET hit or the coiled-coil prediction is taken as reference.

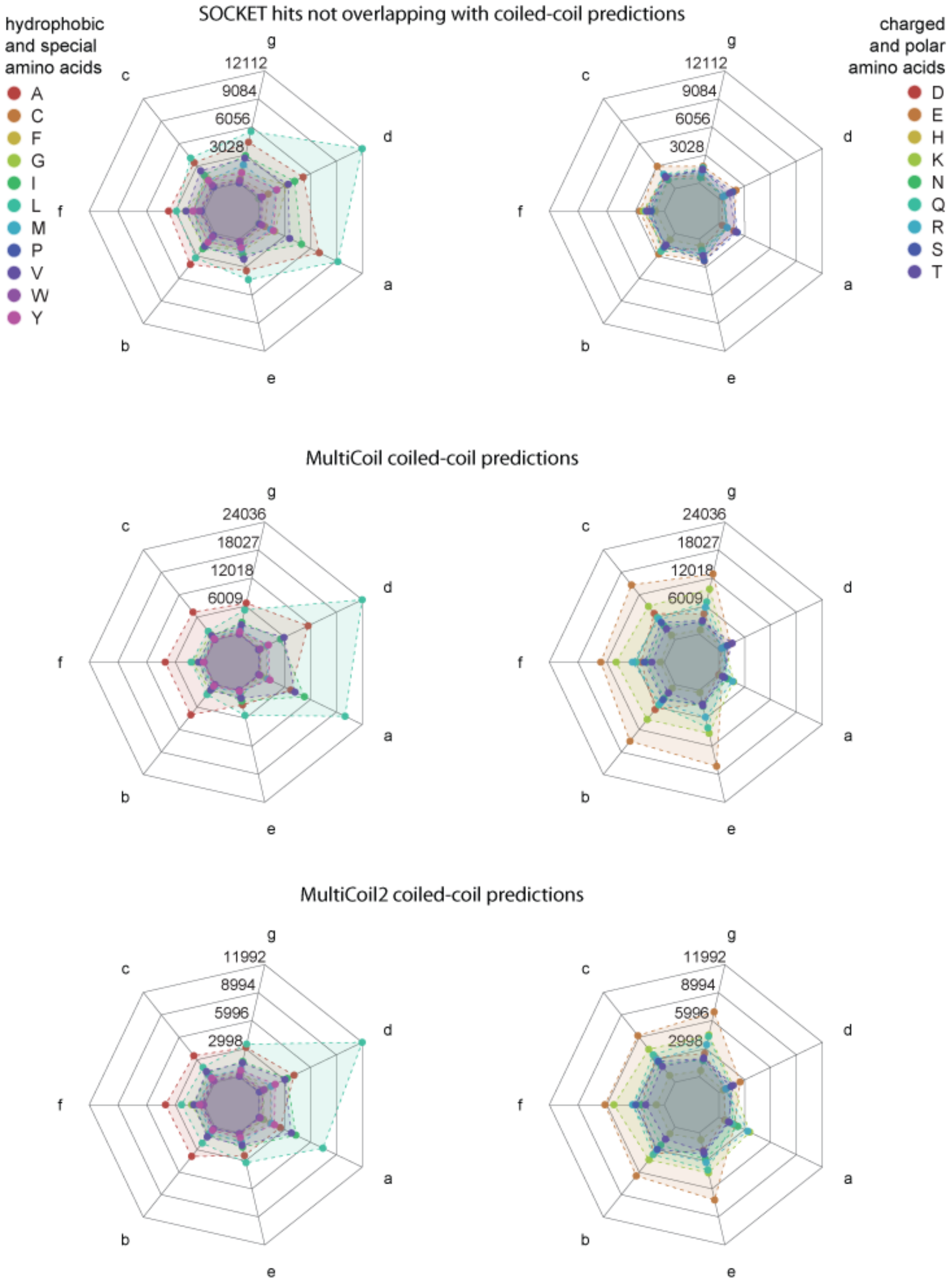
SOCKET hits not overlapping with coiled-coil predictions
hydrophobic and special amino acids
A
C
F
G
I
L
M
P
V
W
Y
charged and polar amino acids
D
E
H
K
N
Q
R
S
T
12112
9084
6056
3028
g
d
a
e
b
f
c
MultiCoil coiled-coil predictions
24036
18027
12018
6009
MultiCoil2 coiled-coil predictions
11992
8994
5996
2998

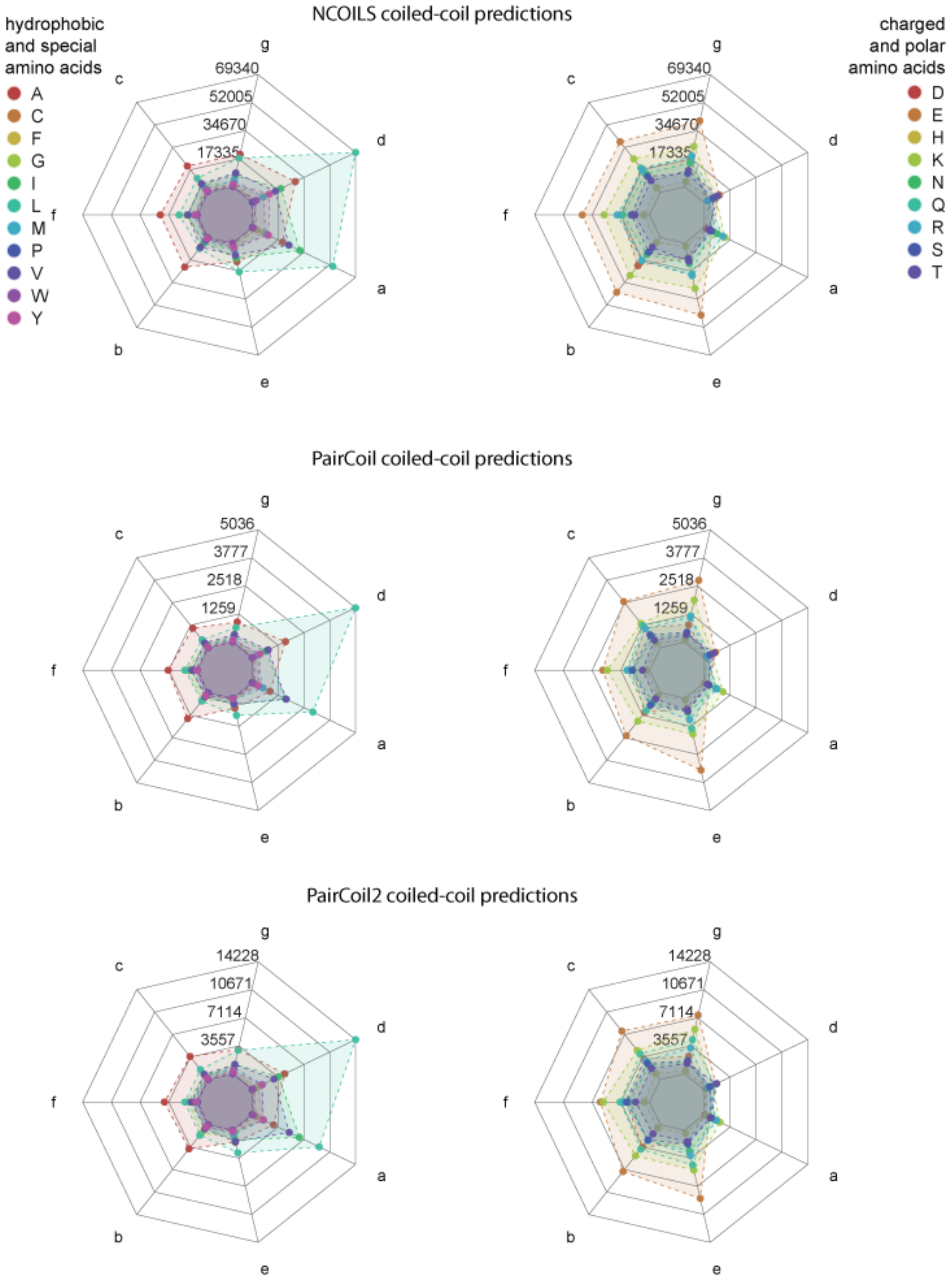


**Figure S3: Amino acid preferences at heptad positions *abcdefg*.** It is well known that hydrophobic amino acids are preferred at the interface between coiled α-helices, at positions *a* and *d*, and that charged and polar amino acids are preferred at the outside, especially at positions *e* and *g*. Because SOCKET not only detects "classical" coiled-coils but interacting α-helices within globular protein structures, the distribution of hydrophobic and charged amino acids is slightly less biased in the latter structures. The heptad patterns of the coiled-coils predictions show strong bias for leucine and isoleucine at the interior positions *a* and *d*, and for glutamate at all other positions, independently of the tool. The letters at the axes denote the heptad register positions. Data values at grid lines refer to amino acid counts at each heptad position over all heptads.

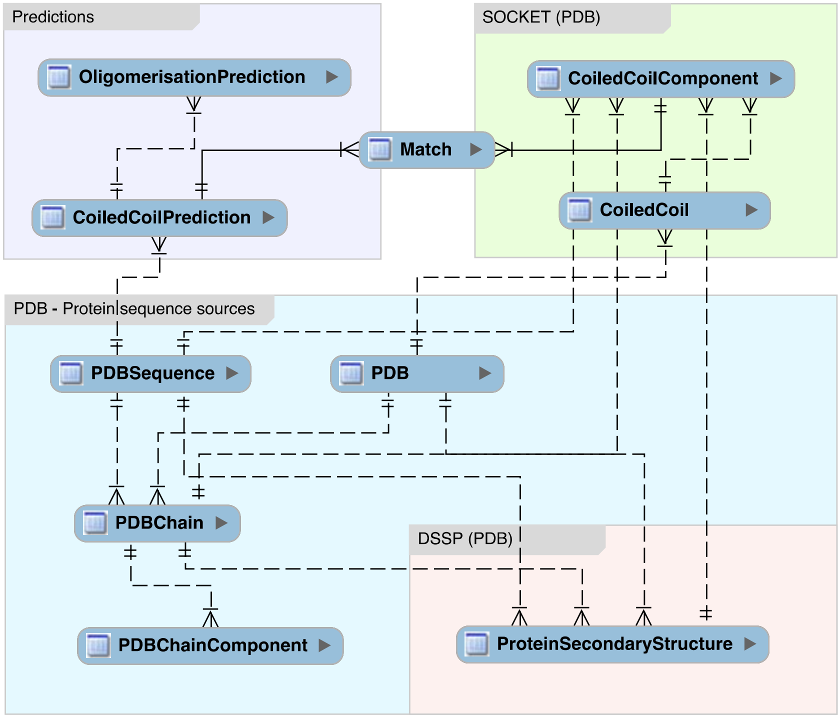


**Figure S4: Database scheme.** Only the table names and connections are shown for clarity.
